# FGFR1 governs iron metabolism via regulating post-translational modification of IRP2 in prostate cancer cells

**DOI:** 10.1101/2022.10.17.512481

**Authors:** Hui Lin, Liuhong Shi, Dongyan Zhou, Shuangya Chen, Ping He, Xiaolu Zheng, Feng Qiu, Yuying Yuan, Shuaijun Lin, Xiaokun Li, Fen Wang, Cong Wang

## Abstract

The acquisition of ectopic fibroblast growth factor receptor 1 (FGFR1) expression is well documented in prostate cancer (PCa) progression. However, how FGFR1 facilitates PCa progression is not fully revealed, although it is known to confer tumor growth advantage and metastasis. Here we report that FGFR1 deletion in DU145 human PCa cells retards the iron metabolism and reduces transferrin receptor 1 (TFR1), which synergistically enhances the anti-cancer effect of iron chelator. FGFR1 and TFR1 are highly expressed in PCa, and FGFR1 overexpression increased TFR1 in PCa cell lines. Furthermore, we first time demonstrate that FGFR1 deletion boosts and shifts the degradation of iron regulatory proteins 2 (IRP2) to downregulate TFR1. Detailed characterization revealed that based on FGFR1 deletion the stability of IRP2 is broken, whose degradation is accelerated, which can be not observed without FGFR1 deletion. In addition, IRP2 overexpression rescue the malignancy degree of DU145 cells. Our results here unravel a novel mechanism by which FGFR1 promotes PCa progression by upregulating iron metabolism, and that the FGFR1/IRP2/TFR1 axis can be a potential target for managing PCa progression.

## Introduction

Iron is an essential element necessary for the basic functions of cells. Normally, iron is bound to transferrin (TF) in the cell membrane. The iron-loaded TF then forms a complex with the transmembrane TF receptor 1 (TFR1) and is internalized by endocytosis (Daniels *et al*, 2006; Hentze *et al*, 2010). In endosomes, members of the metalloreductases function as iron reductase, which reduces Fe^3+^ to Fe^2+^ (Crielaard *et al*, 2017; Zhou *et al*, 2018). Fe2+ is then transported by divalent metal-ion transporter 1 (DMT1), zinc transporter, and iron-related protein 14 (Zrt- and Irt-like protein 14, ZIP14) or ZIP8 from the endosome to the cytoplasm (Brookes *et al*, 2006b; Zhang & Zhang, 2015). The free iron in the cytoplasm constitutes labile iron pool (LIP), which is metabolically active.

Increased total iron content in cancer cells partially resulted from increased TFR1 expression. TFR1 overexpression promotes tumorigenesis and proliferation through increasing LIP in tumor cells (Huang *et al*, 2022; Jung *et al*, 2021; Lodhi *et al*, 2021; Xiao *et al*, 2020). The master regulator of TFR1 is iron regulatory protein 2 (IRP2), which binds to a specific sequence of nucleotides, termed iron responsive elements (IREs) (Pantopoulos, 2004). IRP2 binds the IREs in 3’ untranslated region of TFR1 mRNA, which stabilizes it and promotes the translation to proteins. IRP2 does not sense iron. However, iron regulates IRP2 degradation through the iron regulated FBXL5 that is a E3 ubiquitin ligase for IRP2 degradation (Rouault & Maio, 2020; Terzi *et al*, 2021; Wang *et al*, 2020). Thus, iron-mediated IRP2 degradation controls TFR1 expression and therefore, iron content in the cells.

Prostate cancer (PCa) is the most frequently diagnosed malignancy and the second leading cause of cancer death in men (Torre *et al*, 2015). Organ-confined PCa can be effectively treated by androgen deprivation. However, such treatment inevitably leads to the recurrence of cancer, which generally is metastatic and androgen deprivation insensitive (Isbarn *et al*, 2009). Ectopic expression of fibroblast growth factor (FGF) and FGF receptor (FGFR) is frequently found associated with a variety of human cancers, including colon, lung, bladder, and prostate cancers (Wesche *et al*, 2011). Consistently, deletion of *FGFR1* in prostate epithelial cells impedes the initiation and progression of PCa in both in vitro and in vivo (Corn *et al*, 2013; Yang *et al*, 2013). However, how ectopically expressed FGFR1 contributes to PCa initiation and progression is not fully unraveled. Understanding the molecular mechanism by which FGFR1 contributes to PCa progression will facilitate the development of novel approaches for treating this deadly disease.

Malignant transformation of prostate epithelial cells is associated with increased intracellular iron content (Choi *et al*, 2008; Ornstein & Zacharski, 2007). However, the underlying mechanism is not elucidated. Whether and how ectopic FGFR1 contributes to dysregulated iron metabolism also remains unknown. To address these issues, we first characterized the expression patterns of iron metabolism-related molecules in human PCa tissues and PCa cells and then delineated how FGFR1 regulated iron metabolism. Our results suggest that IRP2/TFR1 and LIP are involved in FGFR1-mediated PCa growth and progression.

## Results

### TRAMP PCa expresses ectopic FGFR1, and exhibits increased the iron abundance

Iron metabolism is closely related to cancer progression (Kerins & Ooi, 2018; Luscieti *et al*, 2017; Wang *et al*, 2017). Disorders of iron metabolism, especially excessive iron acquisition and retention, induces tumor initiation and progression (Jung *et al*, 2017; Schoenfeld *et al*, 2017). To investigate FGFR1 and iron content of prostate and PCa in the TRAMP model, prostate and PCa tissues from various ages of mice were collected for detailed characterization. No apparent differences were observed in the morphology of prostate between wild and TRAMP group, either 12-week-old or 22-week-old (Fig. 1A). However, the expression of FGFR1 and iron content were significantly increased in PCa of 22 weeks TRAMP mice compared with PCa tissues from 12-week-old mice besides the distortion of gland structure, which were not examined in wild type mice at different age (Fig 1B, C&D). Consistently, immunostaining exhibited TFR1 was dramatically expanded in TRAMP mice along with the PCa progression (Fig. 1E). In addition, Perl’s staining of human PCa tissues also showed that poorly PCa had higher iron than that in well differentiated PCa (Fig. 1F).

**Figure 1.**
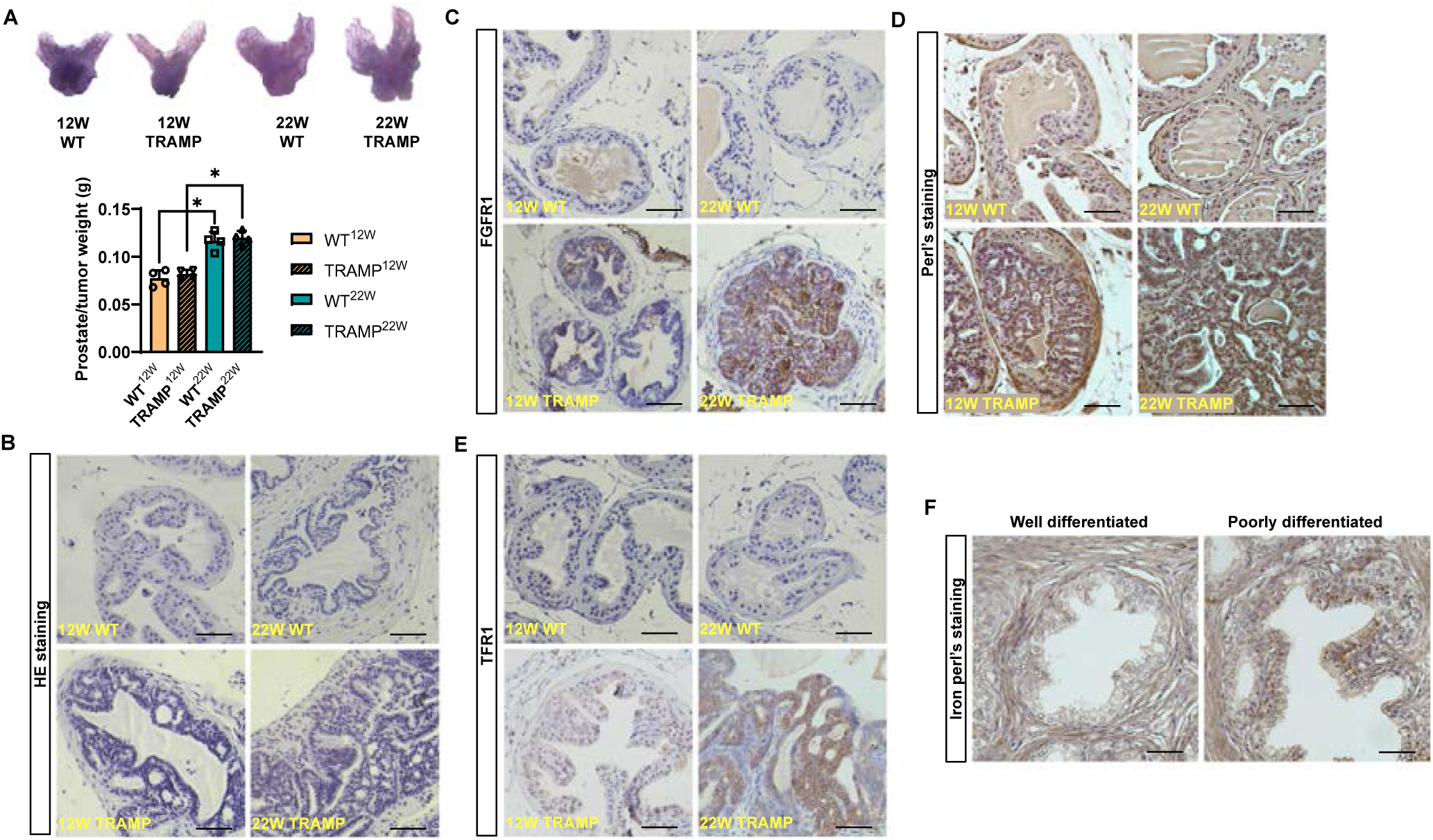
TRAMP PCa expressed ectopic FGFR1 and exhibited the increased iron abundance. **A.** The morphology of prostate in WT and TRAMP mice dissected at the age of 12 and 22 week is observed in a light microscope. Below is the analysis of prostate/tumor weight. (n = 4 mice for each group) **B.** Representative image of H&E staining. Scale bars: 50μm. **C.** Immunohistochemistry staining showing the expression of FGFR1 (brown). Hematoxylin for the counterstaining of nuclear. Scale bars: 50μm. **D.** Representative image of Perl’s staining enhanced with DAB. Scale bars: 50μm. **E.** Immunohistochemistry staining showed the expression of TFR1 (brown). Scale bars: 50μm. **F.** Representative Perl’s staining enhanced with DAB of well differentiated and poor differentiated human PCa tissues. Scale bars: 50μm. Data information: In A, data are presented as mean ± SD. 12W, mice at the age of 12 week. 22W, mice at the age of 12 week. WT, wildtype mice. TRAMP, TRAMP mice. *, *P* < 0.05.

### FGFR1 deletion potentiated the inhibitory effect of DFO on PCa cells

To determine whether ectopic FGFR1 expression was associated with iron metabolism in PCa, we generated FGFR1 null DU145 cells (DU145^ΔR1^ cells). The size of labile iron pool (LIP) of DU145^ΔR1^ cells was significantly decreased compared with control cells (Fig. 2A). The decreased LIP in DU145^ΔR1^ cells prompted us to assess expression of TFR1, and DMT1 in FGFR1 null DU145^ΔR1^ cells. The results showed that the expression of TFR1 and DMT1 was reduced at both mRNA and protein levels in DU145^ΔR1^ cells (Fig. 2B&C), suggesting that ablation of FGFR1 reduced LIP in PCa.

**Figure 2.**
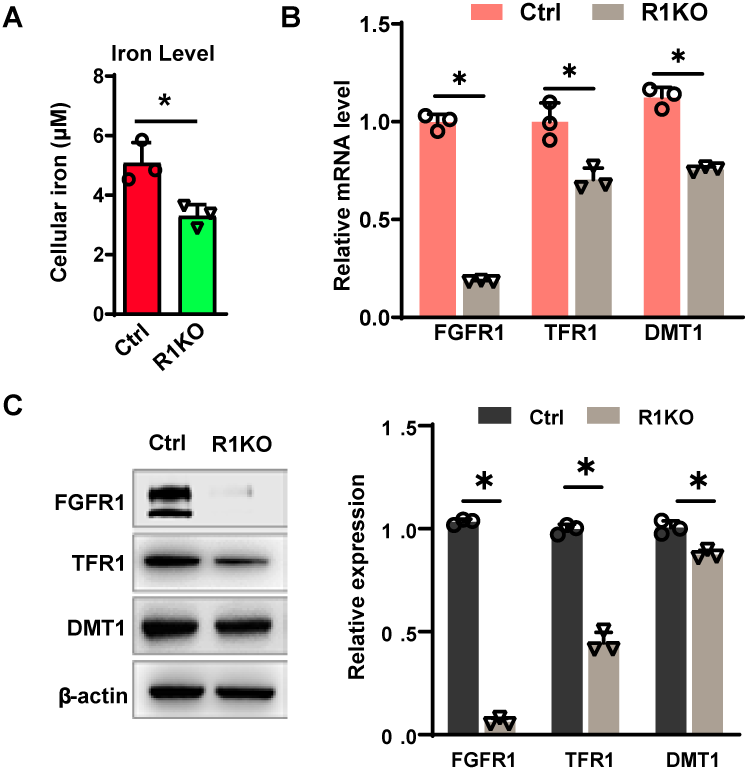
FGFR1 deletion retarded the turnover of iron in PCa cells. **A.** Cellular iron content of DU145 and DU145^ΔR1^ cells detected by iron assay kit. **B.** Relative mRNA expression of iron transport proteins in DU145 and DU145^ΔR1^ cells. **C.** Western blotting analysis of the indicated proteins and the quantitative analyses with Image J. β-actin was used as a loading control. Data information: In (A-C), data are presented as mean ± SD. Ctrl, DU145. R1KO, DU145^ΔR1^. *, *P* < 0.05.

To determine whether ablation of FGFR1 sensitized the cells to iron deficiency, the cells were treated with DFO, an iron-chelating agent, and then subjected to cell viability analysis. The CCK-8 analysis revealed that DFO inhibited cell viability more potently in DU145^ΔR1^ cell than in parental cells (Fig. 3A). Furthermore, EdU incorporation assays showed that DFO suppressed cell proliferation more significantly in DU145^ΔR1^ cells than in parental cells (Fig. 3B). Consistently, expression of PCNA and cMyc, the cell proliferation related protein was lower in DU145^ΔR1^ than in parental cells upon DFO treatment (Fig. 3C).

**Figure 3.**
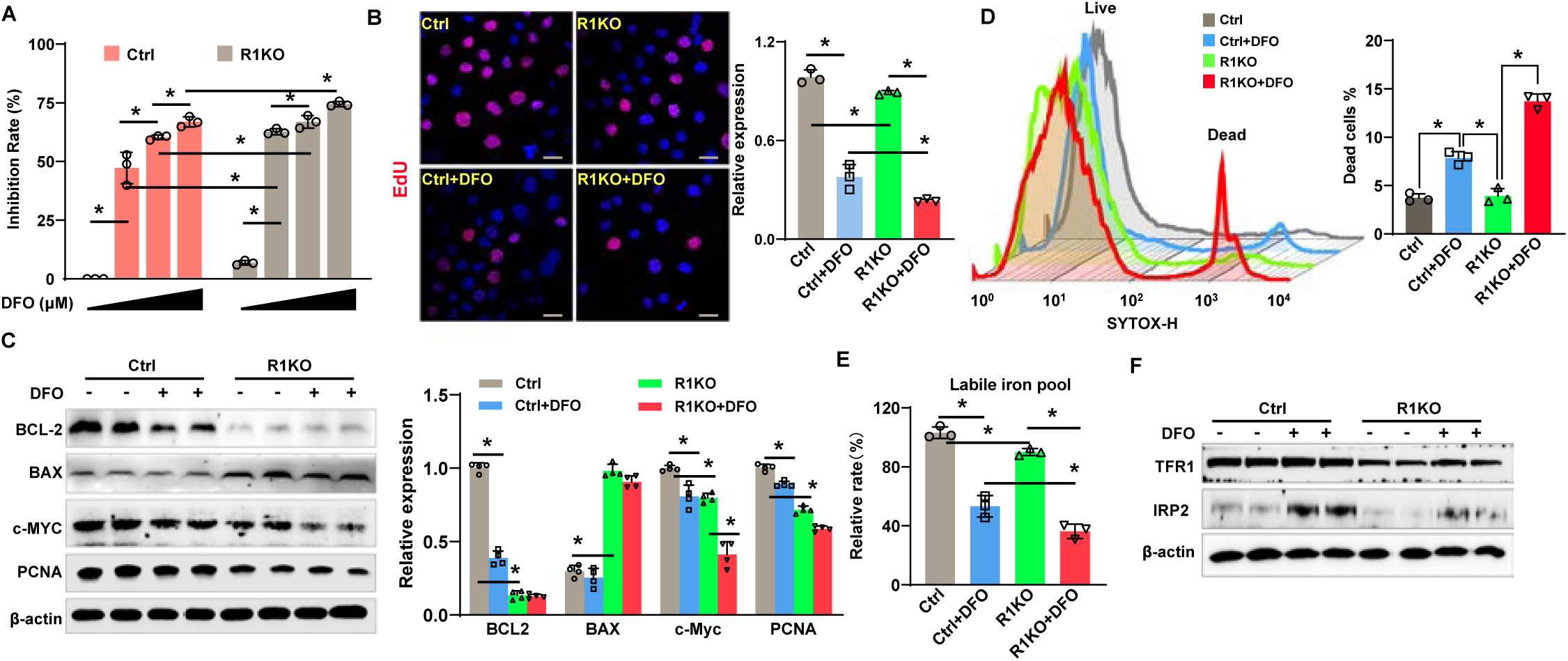
FGFR1 deletion potentiated the inhibitory effect of DFO on PCa cells. **A.** CCK8 analysis was performed to analyze viability of DU145 and DU145^ΔR1^cells treated with DFO at different concentration (0 μM, 20 μM, 50 μM and 100 μM) for 24 hours. **B.** EdU incorporation assay was used to analyze the proliferation of R1KO cells treating with 20 μM DFO, the ratio of EdU positive cells were calculated in three random areas by Image J software, DAPI (blue) for the counterstaining of nuclear. Scale bars: 30μm. **C.** Western blotting analysis of the indicated proteins for proliferation and apoptosis, the intensity was quantitated with Image J. β-actin was used as a loading control. **D.** SYTOX Green was used to detect the survival of DU145 and DU145^ΔR1^cells treated with DFO for 24h, the dead cells were observed by FCM, the ratio of SYTOX positive cells were calculated by three different tests. **E.** Labile iron pool of DU145 and DU145^ΔR1^cells treated with DFO were detected with Calcein AM. **F.** Cells treated with DFO for 24h were lysed for Western blot analysis with indicated antibodies and the intensity was quantitated with Image J. β-actin was used as a loading control. Data information: data are presented as mean ± SD. Ctrl, DU145. R1KO, DU145^ΔR1^. *, *P* < 0.05.

To determine how ablation of ectopic FGFR1 contributed to DFO-induced apoptosis, SYTOX Green labeling was used to quantitate the percentage of apoptotic. Although there was no noticeable difference of apoptotic cells between DU145^ΔR1^ and parental cells, upon DFO treatment, DU145^ΔR1^ cells had significantly more apoptotic cells than the parental cells (Fig. 3D). Western blotting also showed that BCL2 expression in DU145^ΔR1^ cells was downregulated, and BAX was upregulated (Fig. 3C).

Interestingly, LIP in DFO treated DU145^ΔR1^ cells was lower than in controlled cells under the same treatment (Fig. 3E), suggesting that depletion of ectopic FGFR1 signaling exhibited synergistic effect with DFO. Moreover, treating DU145 cells with DFO significantly increased expression IRP2 (Fig. 3F), indicating that low LIP induced feedback upregulation of IRP2. However, such feedback regulation was diminished in DU145^ΔR1^ cells, suggesting that FGFR1 was required for the feedback regulation of IRP2.

### Ectopic expressed FGFR1 enhanced the expression of TFR1 in PCa progression

Ablation of FGFR1 reduced TFR1 expression in DU145 cells (Fig. 3F). Previous reports indicated that the expression of TFR1 was closely related to the development of PCa, liver cancer, and colon cancer (Beckman *et al*, 2000; Habashy *et al*, 2010; Pizzamiglio *et al*, 2017). Therefore, we assessed the association of TFR1 expression with Gleason scores in the TCGA database. The result showed that expression of TFR1 at the mRNA level was positively associated with Gleason scores (Fig. 4A). In addition, the expression of FGFR1 at the mRNA level was also positively associated with that of TFR1 in the same TCGA database (Fig. 4B). Immunostaining demonstrated that poorly differentiated PCa expressed higher FGFR1 and TFR1 than the well differentiated PCa (Fig. 4C). Consistent with the data that TFR1 expression in DU145^ΔR1^ cells was lower than in control cells (Fig. 4D), reinstating FGFR1 expression in DU145^ΔR1^ cells rescued the decreased TFR1 and LIP (Fig. 4E&F). In addition, the expression of FGFR1 in LNCaP and C4-2 cells was below the detection limited, forced expression of FGFR1 in LNCaP and C4-2 cells significantly increased the expression of TFR1 (Fig. 4G&H). Consistently, forced expression of FGFR1 also increased LIP in these cells (Fig 4I). The results indicated that ectopic expressed FGFR1 promotes TFR1 expression and therefore, increases LIP in PCa.

**Figure 4.**
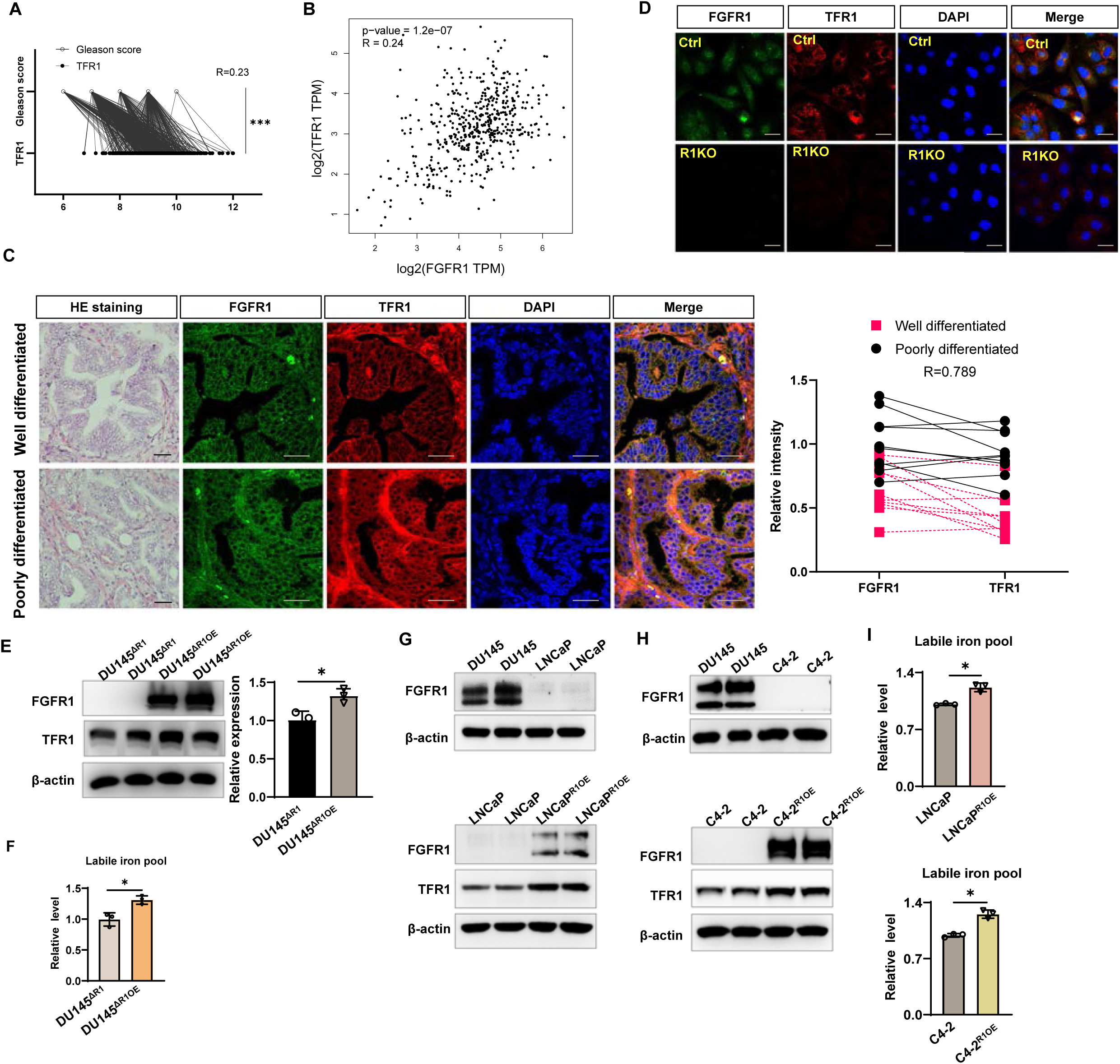
Ectopic expressed FGFR1 enhanced the expression of TFR1 in PCa progression. **A.** Relationship between Gleason score and the mRNA level of TFR1 of prostate cancer in TCGA database. **B.** Correlation analysis between the mRNA level of TFR1 and FGFR1 of prostate cancer in TCGA database. **C.** HE staining and double immunofluorescence staining with FGFR1 (green) and TFR1 (red) between well and poorly differentiated human PCa tissues. Right panel is the quantitative analysis by Image J. Scale bars: 30μm. **D.** Double immunofluorescence staining with FGFR1 (green) and TFR1 (red) in DU145 and DU145^ΔR1^ cells. Scale bars: 30μm. **E.** Western blotting analysis of the indicated proteins and the quantitative analyses of TFR1 with Image J. β-actin was used as a loading control. **F.** Calcein AM was used to detect the labile iron pool of DU145 and DU145^ΔR1^cells. **G.** Western blot showed the protein expression of FGFR1 and TFR1 in different PCa cell lines and the quantitative analyses of TFR1 in indicated PCa cells with Image J. β-actin was used as a loading control. **H.** Labile iron pool were detected with Calcein AM. Data information: In (E-H), data are presented as mean ± SD. Ctrl, DU145; R1KO, DU145^ΔR1^; DU145^ΔR1OE^, FGFR1 overexpressed in DU145^ΔR1^; LNCaP ^R1OE^/C4-2 ^R1OE^, FGFR1 overexpressed in LNCaP/C4-2. ***, *P* < 0.001. *, *P* < 0.05.

### FGFR1 deletion accelerated the degradation of IRP2

To determine how FGFR1 deletion affected expression TFR1, quantitative RT-PCR analyses was employed to assess the expression of IRP1, IRP2, c-MYC and P53, the main upstream regulatory factors of TFR1 (Henderson *et al*, 1996; Liu *et al*, 2019; O’Donnell *et al*, 2006; Zhang *et al*, 2017). The data showed that expression of IRP1 and c-MYC were reduced while that of IRP2 and P53 were increased in DU145^ΔR1^ cells compared with control cells (Fig. 5A). Western blotting analysis also showed that expression of IRP1, IRP2, and c-MYC was reduced while that of P53 was increased (Fig. 5B). Interestingly, the expression of IRP2 was increased at the mRNA level but reduced at the protein level, suggesting that FGFR1 regulated IRP2 expression at the protein level. In addition, forced expression of FGFR1 in C4-2 cells downregulated IRP2 at the mRNA level and upregulated IRP2 at protein level (Fig EV1 A&B). To determine whether FGFR1 suppresses IRP2 degradation, the cells were treated with cycloheximide (CHX) to suppress protein synthesis, and IRP2 proteins level was determined. The results clearly showed that the abundance of IRP2 decreased more rapidly in DU145^ΔR1^ cells than in parental cells (Fig. 5C), suggesting that FGFR1 signaling suppressed IRP2 degradation. Consistently, forced expression of FGFR1 delays IRP2 expression in C4-2 cells (Fig EV1 C).

**Figure 5.**
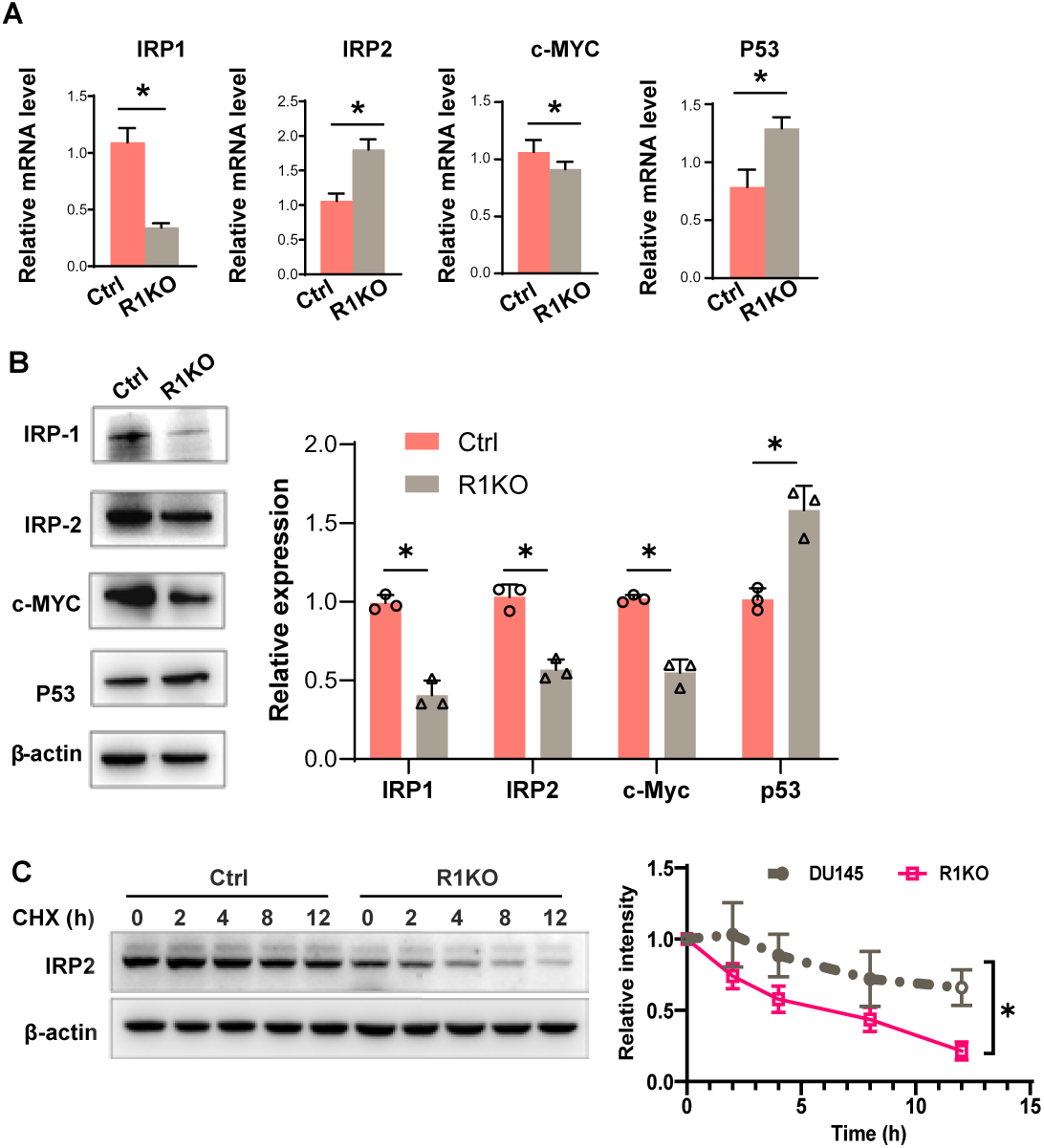
FGFR1 deletion accelerated the degradation of IRP2. **A.** Real-time Q-PCR was performed to analyze relative mRNA expression of IRP1/IRP2/c-MYC/P53 in DU145 and DU145^ΔR1^cells. **B.** Western blotting analysis of the indicated proteins of DU145 and DU145^ΔR1^ cells and the quantitative analyses with Image J. β-actin was used as a loading control. **C.** Western blotting showing the expression of IRP2 in DU145 and DU145^ΔR1^ cells treated with CHX (100μg/ml) for 0 h, 2 h, 4 h, 8 h, and 12 h. Bottom panel is the quantitative analysis by Image J. Data information: In (A-C), data are presented as mean ± SD. Ctrl, DU145. R1KO, DU145^ΔR1^. *, *P* < 0.05.

To determine whether the lysosome-autophagy pathway or ubiquitin-proteasome pathway was involved in IRP2 degradation, the cells were treated with the inhibitors for the lysosome-autophagy or ubiquitylation and then subjected to the same analyses. The results showed that the degradation of IRP2 in DU145 cells was suppressed by proteasome inhibitor MG132, but not by autophagy inhibitor chloroquine (CQ) (Fig. 6A). However, in DU145^ΔR1^ cells, it was suppressed by CQ, but not by MG132 (Fig. 6B), suggesting that IRP2 degradation in control DU145 cells was largely mediated by the ubiquitination-proteosome pathway and in DU145^ΔR1^ cells was mediated by autophagy pathways. The results were in consistence with our previous notion that FGFR1 suppresses autophagy via the mTOR pathways (Lin *et al*, 2011; Liu *et al*, 2018).

**Figure 6.**
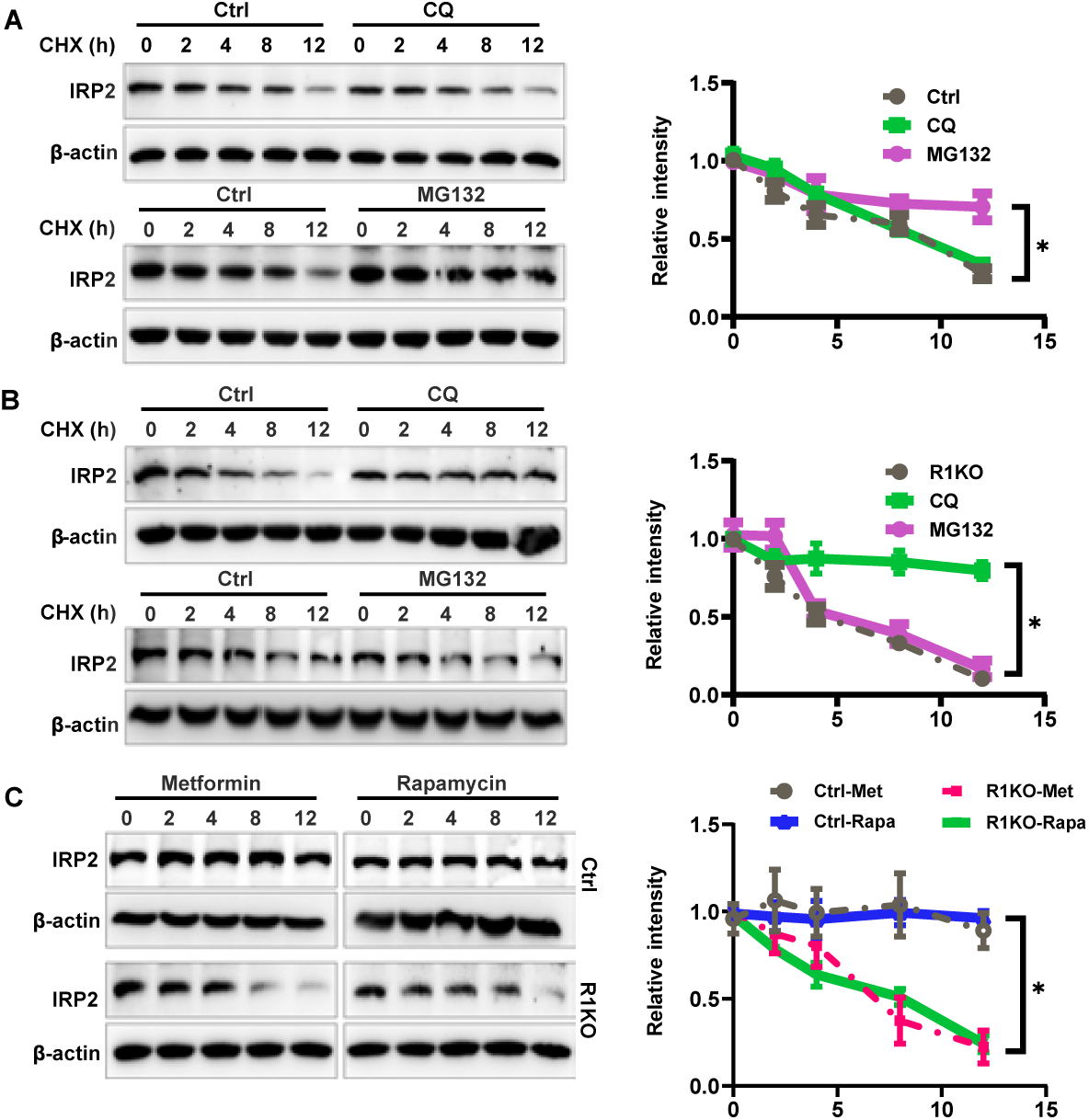
FGFR1 deletion altered the main degradation pathways of IRP2. **A-B.** DU145 (A) and DU145^ΔR1^ (B) cells treated with CHX (100μg/ml) and CQ (50μmol/ml) or MG132 (10μmol/ml) for 0, 2, 4, 8 or 12 hours, and then subjected to Western blot with IRP2 antibody, β-actin was used as a loading control, right panel showed the quantitation of IRP2 by image J. **C.** DU145 and DU145^ΔR1^ cells treated with Metformin (50μmol/ml) or Rapamycin (10μmol/ml) for indicated time, followed by Western blotting showing the expression of IRP2, right panel is time-dependent line charts quantified by image J. Data information: In (A-C), data are presented as mean ± SD. Ctrl, DU145. R1KO, DU145^ΔR1^. *, *P* < 0.05.

To confirm that the activation of autophagy accelerated the degradation of IRP2 in DU145^ΔR1^ cells, the cells were treated with metformin or rapamycin (autophagy activators), and then subjected to the same analyses. The results showed that the treatment of both metformin and rapamycin significantly increased IRP2 degradation in DU145^ΔR^ cells but not in parental cells (Fig. 6C). The results suggest that IRP2 is degraded through autophagy in DU145^ΔR1^ cells.

FBXL5 is the ubiquitin ligase of IRP2 in the ubiquitin-proteosome degradation pathway. Immunostaining showed that IRP2 and FBXL5 were colocalized in DU145 cells, and the colocalization was diminished in DU145^ΔR1^ cells (Fig. 7A). Western blotting revealed that FBXL5 expression was decreased in DU145^ΔR1^ cells (Fig. 7B). The results indicated that IRP2 degradation in control DU145 cells was mediated by the ubiquitination pathway and deletion of FGFR1 compromised this pathway.

**Figure 7.**
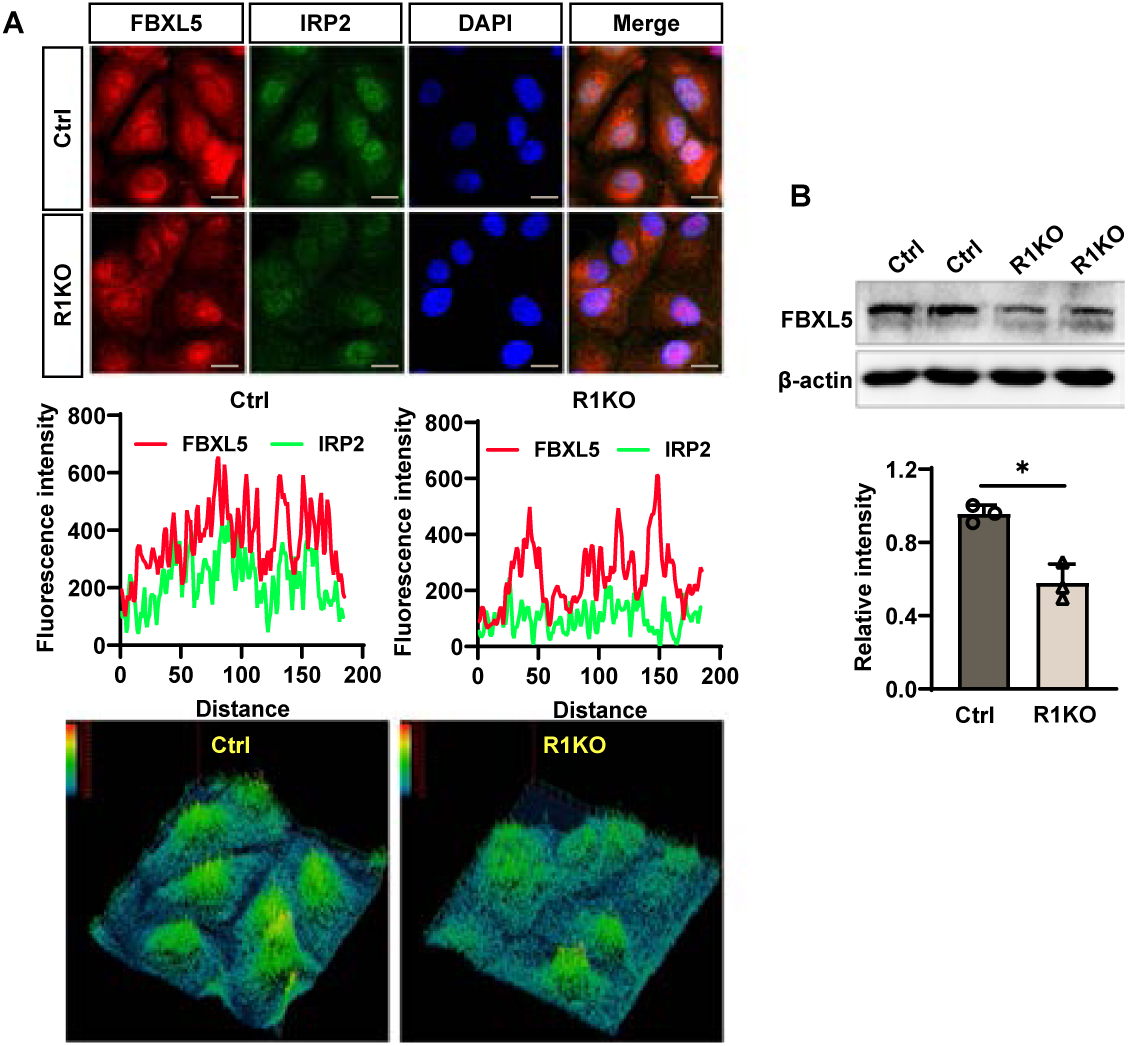
FGFR1 deletion reduced the interactions of FBXL5 and IRP2. **A.** Immunofluorescence staining showing colocalization of IRP2 (green) and FBXL5 (red) in DU145 and DU145^ΔR1^ cells, DAPI for the counterstaining of nucleus. Middle/bottom panel is colocalization analysis and 3D view analysis of colocalization performed by image J software, intensity indication was in the upper left corner of 3D image. Scale bars: 20μm. **B.** Western blotting analysis of FBXL5 and the quantitative analyses with Image J. β-actin was used as a loading control. Data information: In B, data are presented as mean ± SD. Ctrl, DU145. R1KO, DU145^ΔR1^. *, *P* < 0.05.

### Forced expression of IRP2 reinstates the LIP in DU145^ΔR1^ cells

To confirm that reduced IRP2 expression underlie FGFR1 deletion-induced low LIP, DU145^ΔR1^ cells was transfected with IRP2, designated IRP2^OE^. It was clear that the LIP in IRP2^OE^ cells was significantly higher than that in DU145^ΔR1^ cells (Fig. 8A). EdU incorporation assay showed that overexpression of IRP2 reinstated cell proliferation to a level comparable to parental cells (Fig. 8B). Western blotting showed that IRP2 overexpression rescued the expression of TFR1 and FBXL5 (Fig. 8C). Together, the results revealed that ectopic FGFR1 promoted LIP via suppression of ubiquitination-mediated degradation of IRP2.

**Figure 8.**
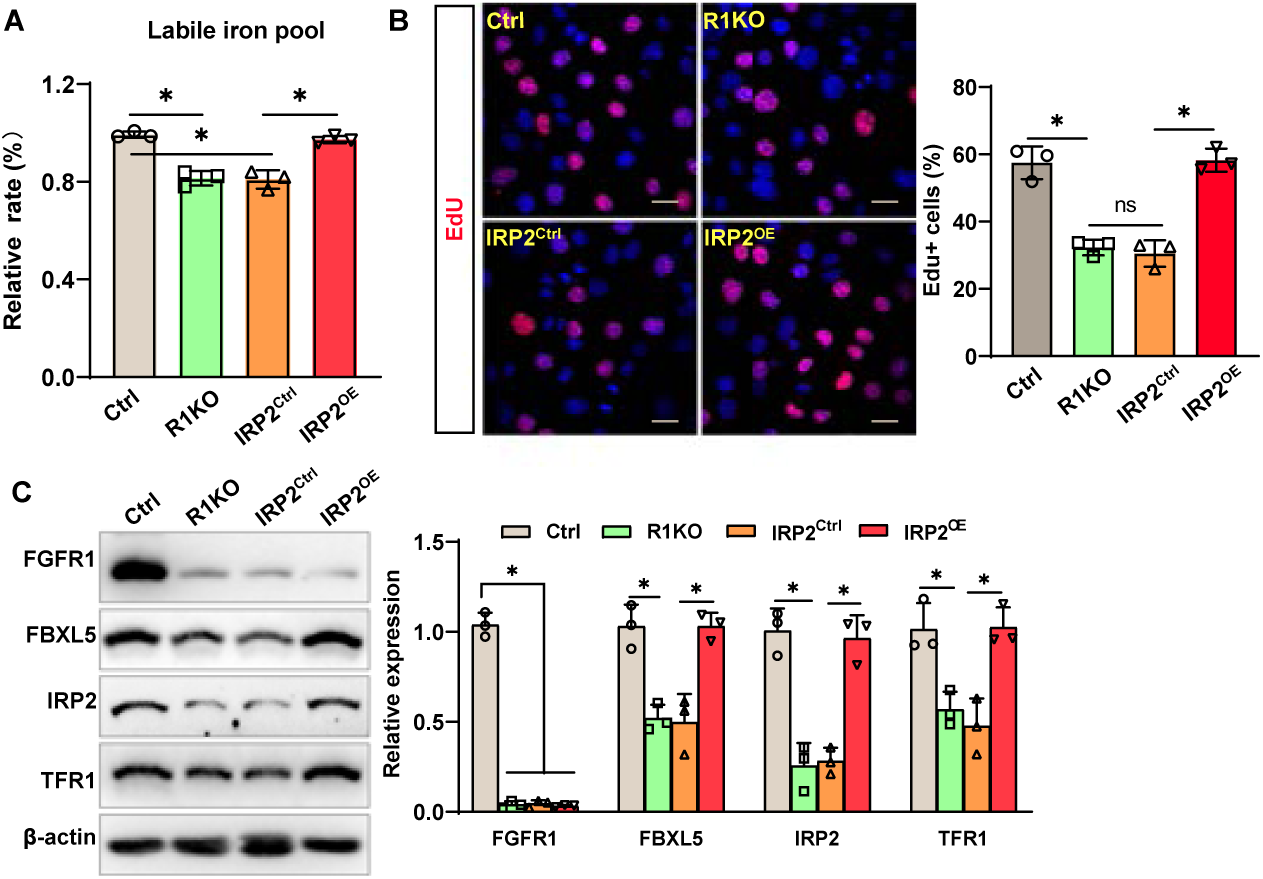
Transfected IRP2 rescued the malignant degree of DU145^ΔR1^ cells. **A.** Statistical analysis of intracellular iron through Calcein AM for indicated cell groups. **B.** EdU incorporation assay was used to analyze the proliferation, the ratio of EdU positive cells were calculated in three random areas by Image J software, DAPI (blue) for the counterstaining of nuclear. Scale bars: 30μm. **C.** Western blotting analysis of indicated proteins and the quantitative analyses with Image J. β-actin was used as a loading control. Data information: In (A-C), data are presented as mean ± SD. Ctrl, DU145; R1KO, DU145^ΔR1^; IRP2^Ctrl^, DU145^ΔR1^ transfected with control plasmid; IRP2^OE^, DU145^ΔR1^ overexpressed with IRP2. *, *P* < 0.05.

**Fig 9.**
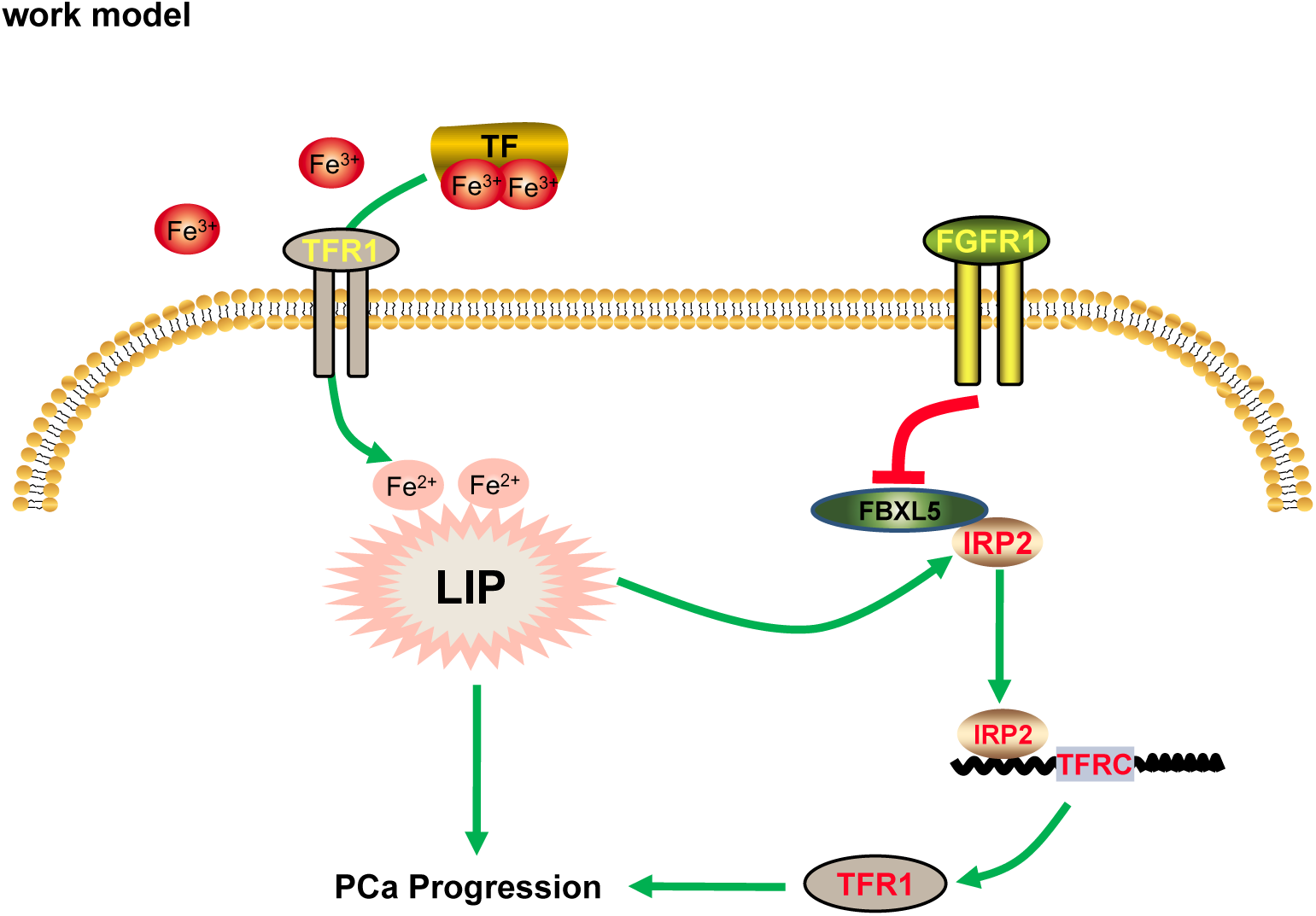
Schematic of FGFR1 governs iron metabolism via regulating post-translational modification of IRP2 in PCa cells. Iron bound to transferrin (TF) uptakes by transferrin receptor 1 (TFR1) on the plasma membrane of cells is the most important way for cancer cells to absorb iron. TFR1 overexpression increases liable iron pool (LIP) in tumor cells to promote tumorigenesis and proliferation. FGFR1 positively regulates the cellular iron content via raising TFR1 expression through inhibiting the degradation of the master regulator, iron regulatory proteins 2 (IRP2), by enhanced FBXL5 activity, which leading to accelerates PCa progression.

## Discussion

Abnormal iron metabolism is emerging as a key metabolic hallmark closely associated with PCa initiation and progression (Beshara *et al*, 1997; Lee & Means, 1995; Shen *et al*, 2018). Also, there is extensive evidence that ectopic FGF/FGFR1 signaling is a contributing factor in PCa development and progression (Abate-Shen & Shen, 2007; Devilard *et al*, 2006; Giri *et al*, 1999; Liu *et al*., 2018; Ozen *et al*, 2001; Taylor *et al*, 2010; Wang *et al*, 2004b). In the present study, we demonstrate that ectopic FGFR1 alters iron metabolism by regulating TFR1 though IRP2 degradation in the iron uptake. FGFR1 deletion in DU145 cells retards the activity of iron metabolism to inhibit cell proliferation and survival. Furthermore, FGFR1 deletion synergistically enhances the effects of the chelator DFO on pro-apoptosis and anti-proliferation. Our data demonstrate that ectopic expression of FGFR1 promotes TFR1 expression through stabilizing IRP2 to increase intracellular iron content. The study first time reports the mechanism of how FGFR1 signaling regulates the iron metabolism in PCa.

Excessive expression of TFR1 and increased iron metabolites are previously reported occurring in a variety of human tumors, including breast, lung, colorectal, and prostate cancers (Brookes *et al*, 2006a; Johnson *et al*, 2014; Jung *et al*., 2021). It has been recently discovered that excessive TFR1 expression becomes a new prognostic factor in various cancers (Johnson *et al*., 2014; Wei *et al*, 2021; Wu *et al*, 2019). Moreover, depleting TFR1 expression has the potential for suppression of tumor progression (Torti & Torti, 2013). However, the mechanism by which FGFR1 regulates iron metabolism is poorly understood. Our data show that elevated expression of FGFR1, TFR1, and iron content are associated with PCa progression in the TRAMP mouse model. FGFR1 deletion decreased TFR1 expression and reduced intracellular LIP. Together, these findings reveal that ectopic FGFR1 expression promotes iron metabolism, and therefore, promotes cell proliferation and survival in PCa cells.

DFO is a commercially available iron chelator that suppresses growth of multiple types of cancers (Komoto *et al*, 2021; Lang *et al*, 2019; Sandoval-Acuña *et al*, 2021). Although DFO has anti-proliferation activity, it is not suitable for cancer treatment, largely due to its short plasma half-life (Blatt, 1994; Selig *et al*, 1998). Herein, we reported that depletion of FGFR1 signaling synergized with DFO in reduction of LIP. Since reduction of LIP sensitizes cells to anti-cancer drugs, it is likely that the combination of FGFR1 kinase inhibitor and DFO can be used to enhance the efficacy of anticancer drugs for PCa.

The iron contents in PCa cells are higher that the adjacent non-cancerous prostate tissues (Guntupalli *et al*, 2007; Sarafanov *et al*, 2011). TFR1 is the major receptor for cellular iron import, the expression of which is significantly elevated in PCa, which has the potential to serve as a marker of PCa diagnosis and prognosis (Keer *et al*, 1990; Zapała *et al*, 2021). In the present results, we found that FGFR1 increased expression of TFR1, depletion of FGFR1 downregulated TFR1 expression, is obviously decreased. Since no TFR1 inhibitors are available, suppression TFR1 expression with the current FDA approved small molecule FGFR1 inhibitors shall be one option to be explored.

IRP2 is a key regulator for iron metabolism, which binds to the IREs in the non-coding region of TFR1 mRNA and prevents its degradation and is a key regulator of iron metabolism (Guo *et al*, 1995; Moroishi *et al*, 2011). Recent reports show that IRP2 degradation is mainly mediated by the ubiquitination. There are various pathways for IRP2 degradation, including iron-dependent and iron-independent pathways (Elton *et al*, 2015; Ishikawa *et al*, 2005; Wang *et al*, 2004a). FBXL5 is an ubiquitin E3 ligase for IRP2 that contributes to iron-dependent degradation of IRP2 (Moroishi *et al*., 2011). However, the iron-independent pathway is not clear. Here we reported that FGFR1 suppressed the non-FBXL5 dependent degradation of IRP2 in PCa, which upregulated the expression of TFR1, resulting in increased LIP in PCa. Depletion of FGFR1 downregulated LIP in PCa, which was rescued by forced expression of IRP2. Thus, inhibiting the FGFR1/IRP2/TFR1 signaling axis has the potential to serve as an effective therapeutic strategy for high iron metabolism cancers, including PCa.

In summary, we demonstrate that ectopic FGFR1 regulates iron metabolism by stabilizing IRP2, and therefore, upregulating TFR1 expression. Depletion of FGFR1 blocks the iron import and reduce LIP, therefore, inhibits cell proliferation and survival. Together with our previous reports that ectopic FGFR1 promotes PCa progression and metastasis, our findings herein unravel a novel mechanism by which ectopic FGFR1 promotes PCa progression and suggest that the FGFR1-iron metabolism pathway can be a novel target to treat currently uncurable castrate-resistant PCa.

## Materials and methods

### Animals and sample preparation

Wildtype male C57BL/6 mice and female hemizygous for the transgenic adenocarcinoma of the mouse prostate (TRAMP, RRID: IMSR_JAX:003135) transgene line, probasin (PB) Tag 8247NG, were purchased from the Jackson Laboratory (Bar Harbor, ME). The male mice were bred on the same genetic background and housed at Laboratory Animal Resources Center of Wenzhou Medical University, maintained under 12 hours light and dark cycles, and provided ad libitum access to food and water. The F1 offspring were genotyped by PCR. Male hemizygous TRAMP mice and male wildtype mice were used. After weaning, mice were randomly divided and housed in the animal facility. At the age of 12 and 22 weeks, mice were sacrificed, and prostate/prostate cancer tissues were collected for analysis.

### Immunostaining assay

For tissue immunofluorescence staining, the sections were deparaffinized, rehydrated, and antigen retrieved. After blocking with 3% bovine serum albumin (BSA, beyotime, China) at room temperature for 30 minutes, the sections were incubated with mixture of primary antibodies overnight at 4℃. After a brief wash, the sections were incubated with mixture of FITC- and Cy5-conjugated secondary antibody (Jackson ImmunoResearch, United States) for 1 hour and DAPI (Cell Signaling Technology, United States) for 5 minutes.

For cell immunofluorescence staining, cells were fixed with 4% paraformaldehyde for 15 minutes, permeabilized with 0.3% Triton X-100/PBS (v/v) for 15 minutes, blocked with 3% BSA at room temperature for 30 minutes, then stained with primary antibodies overnight at 4℃. After a brief wash, the slides were incubated with mixture of secondary antibody for 1 hour and then counterstained with DAPI. Images were taken with a laser scanning confocal microscope (Leica TCS SP8, United States) in three randomly selected areas.

### Prussian blue staining

Prussian blue stain enhanced with DAB (Solarbio, China) was used to detect the presence of iron. Briefly, 5-µm paraffin sections on slides (HistoCore, Leica Biosystems, Germany) were immersed in xylene for 10 minutes. Repeated the step once again in fresh xylene for 10 minutes. Next, the sections were rehydrated by sequentially incubating with 100%, 95%, 80%, and 60% ethanol for 5 minutes each and then rinsed with distilled water three times for 3 minutes each. The sections were incubated for 20 minutes each with Prussian blue, incubation solution, and enhanced solution in sequence. Then the sections were incubated in hematoxylin solution for 40 seconds, followed by a wash with distilled water. Labeled cells were examined under a light microscope (ECLIPSE Ni-E, Nikon, Japan) to determine intracellular iron oxide distribution.

### Cell culture

DU145 cells (ATCC: HTB-81, RRID: CVCL_0105, Manassas, VA, United States) were cultured in Dulbecco’s modified Eagle’s medium (DMEM, Gibco, United States), and LNCaP (ATCC: CRL-1740, RRID: CVCL_1379) and C4-2 (ATCC: CRL-3314, RRID: CVCL_4782) cells were cultured in Roswell Park Memorial Institute (RPMI) 1640 Medium (RPMI1640, Gibco) both supplemented with 10% fetal bovine serum (FBS, Gibco), 100 units/ml penicillin and 100 μg/ml streptomycin in 5% CO_2_ incubators. All the cell lines were tested for mycoplasma contamination routinely every 1 months. All the experiments were performed with cells below the 20th passage and all parental cell lines have been authenticated using short tandem repeat profiling by Genetic Testing Biotechnology Corporation (Suzhou, China) within the last 2 years.

For treatments of cells, iron-chelating agent Deferoxamine (DFO, Sigma-Aldrich, United States), protein synthesis inhibitor Cycloheximide (CHX, Sigma-Aldrich, United States), Proteasome inhibitor (R)-MG132 (Merck, Germany), Lysosomal inhibitors Chloroquine (CQ, Sigma-Aldrich, United States), autophagy inducer Metformin (Met, Aladdin, China), and autophagy inducer Rapamycin (Rap, Aladdin, China) were added to the medium at the indicated concentrations.

### Gene ablation and transfection

The lentivirus-based CRISPR-Cas9 system was used to ablate Fgfr1 alleles in DU145 cells. The sequence of single guide RNA (sgRNA) was AACTTGTTCCGATGGTTATC. Two days after infection with the lentivirus, the virus-containing cells were selected by growing in medium containing 2 μg/ml puromycin. For transient transfections, cells cultured overnight (1×10^5^ cells/well in 6-well plates) were transfected with plasmid and 5μl Lipofectamine 2000 (Thermo Fisher Scientific, United States). The cells were then incubated at 37 °C for 24 h before analysis.

### Measurement of cellular iron level

Cellular iron level was measured by using an iron assay kit (Solarbio, China) according to the manufacturer’s instructions. Briefly, after treatment as indicated, cells were harvested and rapidly homogenized in iron assay buffer. The supernatant was obtained after centrifuging at 16,000g for 10 minutes at 4 °C to remove insoluble material, then added with or without iron reducer and incubated for 30 minutes at 25 °C. Next, the sample was added with an iron probe and incubated at 25 °C for 60 minutes. Finally, the absorbance was measured at 593 nm with a microplate reader (SpectraMax190, MD, United States).

### Quantitative real-time RT-PCR

Total RNA was extracted from cells using the TRIzol RNA isolation reagents (Takara bio, Japan). The first strand cDNA was reverse transcribed from the RNA templates using the GoScript Reverse Transcription system kit (Promega, United States) and Oligo(dT)_15_ primers (Promega, United States) according to the manufacturer’s protocols. Quantified Real-time PCR (qPCR) analyses were carried out using the FastStart Essential DNA Green Master (Roche, Swiss Confederation) as instructed by the manufacturer. The relative abundance of mRNA was calculated using the comparative threshold cycle method and normalized to β-actin as the internal control. Primers are listed in supplementary Table 1.

### Western blotting analysis

Cells were lysed in radioimmune precipitation assay buffer, and the extracted proteins were harvested by centrifugation at 12,000g. Samples containing 30 μg protein were separated on SDS-PAGE and blotted onto polyvinylidene difluoride membranes for Western blot analyses with the indicated antibodies. The primary antibodies are followed: Anti-TFR1 (Cat#sc-32272), anti-DMT1 (1:500), anti-IRP1 (Cat#sc-166022), anti-IRP2 (Cat#sc-33682), anti-GAPDH (Cat#sc-365062), and anti-β-actin (Cat#sc-69879) were all obtained from Santa Cruz Biotechnology. Anti-FGFR1 (Cat#9740), anti-BCL2 (Cat#4223), anti-BAX (Cat#41162), anti-c-MYC (Cat#18583), anti-PCNA (Cat#13110), and anti-P53 (Cat#2527) were obtained from Cell Signaling Technology. Anti-TFR1 (Cat#ab214039), anti-FBXL5 (Cat#ab140175) and anti-FGFR1 (Cat#ab824) were obtained from Abcam. The membranes were washed with TBS with Tween 20 (TBST) buffer to remove nonspecific antibodies and then incubated with horseradish peroxidase-conjugated goat anti-rabbit or anti-mouse IgG (Jackson ImmunoResearch) at room temperature for 1 hour. The specifically bound antibodies were then visualized using ECL-Plus chemiluminescent reagents.

### Cell viability assays

DU145 cells were seeded in 96-well plates (5 × 10^3^ cells per well). The next day, fresh media containing DFO (20μM, 50μM, 100μM), or control (double distilled water) were added, and cells incubated for 24 hours. Cell Counting Kit-8 from MedChemExpress (CCK-8, United States) was used to determine cell viability, as per the manufacturer’s instructions. Cell viability was normalized against the vehicle control, and the data expressed as a percentage of control from three independent experiments done in triplicate.

### EdU incorporation assay

Cells were inoculated 1 × 10^4^ cells per well in 24-well plates, then incubated for 24 hours according to the test requirements by the addition of DFO. Before the termination of cell culture, 5-ethynyl-2’-deoxyuridine (EdU) (10μM, Beyotime, China) was added, and the cells were further incubated at 37°C for 40 minutes. The culture medium was then discarded, and the slide was washed three times with PBS. The cells were fixed with methanol for 10 minutes, and then air dried. After adding the reaction mixture, different fields were randomly chosen to calculate the total number of cells and the number of EdU-positive cells in each vision field.

### Measurement of cell death by FCM

Cell death was measured by staining with SYTOX Green (MedChemExpress, United States) to detect the plasma membrane integrity through FCM according to the manufacturer’s instructions. Briefly, SYTOX Green was excluded from live cells but penetrating dead cells and emitted a green fluorescence that can be quantified by FCM.

### Measurement of cellular iron level

Cellular labile iron was measured according to the manufacturer’s instructions. Briefly, cells were seeded in 96-well plates (5 × 10^2^ cells per well). Cells were incubated with 1 μM Calcein AM for 30 min at 37 °C. Then, cells were washed twice with PBS. The samples were examined by microplate readers (Synergy NEO2, BioTek, United States) followed by incubated with 10μM Pyridoxal isonicotinoyl hydrazone (PIH, MedChemExpress, United States) for 5min. The samples were examined by microplate readers again. The difference between the two fluorescence intensities is the intracellular LIP.

### Correlation analysis of TCGA data

RNA-sequencing expression profiles and corresponding clinical Gleason score for prostate cancer were downloaded from the TCGA dataset (https://portal.gdc.com). The two-index correlation map is realized by correlation analysis and visualized. Pearson’s and Spearman’s correlation analysis to respectively describe the correlation between quantitative variables with or without normal distribution. P values less than 0.01 were considered statistically significant.

### Hematoxylin and Eosin (H&E) staining

Paraffin sections were dewaxed in xylene for 5-10 minutes, switched to fresh xylene and dewaxed for another 5-10 minutes, which were incubated in anhydrous ethanol for 5 min, 90% ethanol for 2 min, 80% ethanol for 2 minutes, 70% ethanol for 2 min, and distilled water for 2 minutes. Sections were stained by hematoxylin staining solution for 5 minutes, immersed in tap water and rinsed off excess staining solution for about 10 minutes, washed again with distilled water, and stained by eosin solution for 30 seconds. After dehydrated, cleared and mounted, sections were observed under a light microscope (ECLIPSE Ni-E, Nikon, Japan) to determine morphological changes.

### Statistical analysis

Statistical analysis was done using GraphPad Prism 8.0.2 software (GraphPad Software, CA). Results were shown as mean ± SD from three or more duplicated tests. The sample size refers to biological repeats, which means repeating using a different biological sample preparation. P-Values among two groups were computed using Student’s t-test. And P-Values among exceeding three groups were computed using one-way ANOVA. Results were considered significant if *P* < 0.05.

## Acknowledgements

This work was supported by the National Natural Science Foundation of China (82173013, 81971894), the Natural Science Foundation of Zhejiang Province of China (LR20H310001, LWY20H300001, LY17H150003), Project of Wenzhou Science & Technology Bureau (Y20210085).

## Author Contributions

Conceptualization: C.W., X.L.; Methodology: H.L., L.S., D.Z., S.C., P.H., X.Z., F.Q., Y. Y., S. L.; Software: H.L., L .S.; Validation: H.L., P.H., S.C., X.Z., F.Q.; Formal analysis: S.C., H.L., P.H.; Investigation: C.W., X.L., F.W.; Resources: F.W., C.W.; Data curation: H.L., D.Z., L.S.; Writing - original draft: H.L., C.W; Writing - review & editing: C.W., F.W.; Project administration: C.W., F.W.; Funding acquisition: D.L., C.W., X.L. The work reported in the paper has been performed by the authors, unless clearly specified in the text.

## Data Availability

This study includes no data deposited in external repositories.

